# Overcoming treatment resistance mediated by the bone marrow vascular niche in acute myeloid leukemia

**DOI:** 10.1101/2025.05.22.655592

**Authors:** Matthew Froid, Sergio Branciamore, Ziang Chen, David Frankhouser, Yu-Hsuan Fu, Jennifer Rangel Ambriz, Le Xuan Troung Nguyen, Jihyun Irizarry, Ya-Huei Kuo, Denis O’Meally, Bin Zhang, Guido Marcucci, Russell Rockne, David Basanta

## Abstract

Acute myeloid leukemia (AML) is a hematologic malignancy originating in the bone marrow, frequently progressing to extramedullary sites. Despite advances in molecularly targeted therapies and hematopoietic stem cell transplantation, clinical outcomes remain suboptimal. Tyrosine kinase inhibitors (TKIs) confer benefit in a subset of AML patients harboring FLT3-ITD mutations, yet relapse and resistance are common. These failures are driven by both intrinsic properties of leukemic stem cells (LSCs)—a quiescent, self-renewing population—and extrinsic cues from the tumor microenvironment. We previously demonstrated that arteriolar endothelial cells (ECs) produce miR-126, which is transferred to LSCs, promoting quiescence, treatment resistance, and niche retention. During disease progression, TNF-α secreted by expanding blasts suppresses EC miR-126 production, enabling LSCs and their progeny to proliferate. Following TKI administration, blast reduction lowers TNF-α levels, restoring EC miR-126 production and enabling LSCs to re-enter quiescence—thereby escaping therapy and facilitating relapse.

To investigate this dynamic, we developed an agent-based computational model of the AML bone marrow microenvironment, parameterized with in vitro and in vivo data. The model captures vascular niche remodeling and the feedback between leukemic populations and endothelial signaling. Simulations reveal that LSC protection mediated by miR-126 can be overcome by combining TKIs with miRisten, a miR-126 inhibitor. When administered on a defined schedule, this combination disrupts the protective niche and enhances LSC eradication. These findings underscore the therapeutic potential of targeting microenvironmental feedback to overcome resistance and prevent AML relapse.

## Introduction

Acute myeloid leukemia (AML) is an aggressive hematologic malignancy driven by genetic mutations, chromosomal aberrations, and epigenetic modifications that promote expansion of partially differentiated leukemic cells (blasts) that ultimately disrupt normal hematopoiesis. Leukemia stem cells (LSCs), a primitive subpopulation of leukemic cells with unlimited self-renewal capacity and high treatment resistance, are responsible for initiation and persistence of maintenance of the disease and treatment failure. Thus, a central strategy for achieving long-term remission and a cure in AML is LSC eradication. Although allogeneic stem cell transplantation can be curative and eliminates LSCs through a graft-versus-leukemia effect, its application is limited by substantial morbidity and mortality, making it unsuitable for elderly or medically unfit patients [1-3].Therefore, the development of safer and more effective therapeutic approaches remains an urgent clinical priority for this disease.

Among the distinct molecular subtypes of AML, more than 20% of cases harbor an internal tandem duplication (ITD) mutation in the FLT3 gene (FLT3-ITD), which encodes a constitutively active mutant tyrosine kinase that drives leukemic growth [4, 5]. The clinical benefit of integrating of tyrosine kinase inhibitors (TKIs) into the treatment of FLT3-ITD AML has been demonstrated in several clinical trials [6-9]. However, TKI resistance emerges not only through mechanisms that are intrinsic to LSCs but also through the functional interplay of LSCs with the bone marrow (BM) tumor microenvironment (i.e., LSC niche) [10-12]. We previously reported that while cytotoxic treatment with TKIs initially induce cytoreduction of FLT3-ITD leukemic blasts, it also allows for arteriolar revascularization of the leukemic BM niche thereby resulting in LSC protection and paradoxically promoting disease relapse [11]. We identified miR-126 as a critical mediator of this dynamic process, as endothelial cell (EC) miR-126 regulates BM arteriolar revascularization supplied and is supplied by these vessels to LSCs, where it governs homeostasis and quiescence.

MicroRNAs (miRNAs) are short non-coding RNAs that regulate gene expression by modulating target messenger RNA levels and regulating protein production. MiR-126 has been reported to contribute to LSC quiescence and treatment resistance [13]. This microRNA is highly produced in arteriolar ECs and exported to LSCs. During LSC-driven disease growth, the myeloblast progeny secretes pro-inflammatory cytokines, such as TNF-α, that suppress EC miR-126 production and in turn reduce the supply to LSCs which exit quiescence and ignite an accelerated leukemia growth. However, upon initiation of cytoreductive therapy, the myeloblasts are killed, and the reduced TNF-α production causes a surge into EC miR-126 production and supply to LSCs. LSCs then retract into a quiescent and treatment resistant state, until the drug is cleared, upon which they restart another cycle of disease growth (i.e., relapse). This biphasic and paradoxical response to therapy which ultimately leads to LSC persistence and disease relapse [11], exemplifies a treatment-related “Janus phenomenon” in AML, where the initial therapeutic benefit of leukemic blast reduction is offset by a detrimental, treatment-induced remodeling of the BM microenvironment and production of EC miR-126 protective of LSCs. To counteract the treatment induced Janus phenomenon, we investigated miRisten, a novel miR-126 inhibitor [12], and hypothesized that optimization of timing and scheduling of miRisten administration in relation to TKI or other cytotoxic therapies may prevent the Janus phenomenon by reducing EC miR-126 production and supply to LSCs [10-12, 14, 15].

Using experimental data, we developed an agent-based model (ABM) to simulate the Janus phenomenon induced by TKIs and explore how different vascular architectures and drug schedules could lead to elimination of LSCs and mitigate the risk of disease relapse. ABMs are computational models that simulate interactions among autonomous agents to better understand system-wide behaviors and outcomes. These stochastic models are built from the bottom up, assigning defined attributes to individual agents and programming them to interact dynamically with their environment and each other. The emergent effects from these interactions often differ from those predicted by individual agent behaviors. Our ABM was designed to replicate this phenomenon and suggests that the Janus effect in AML can be overcome through strategic treatment scheduling, such as pretreatment with miRisten followed by concurrent administration of a TKI and miRisten, or through sequential monotherapy with miRisten followed by a TKI (AC220). These findings not only underscore the pivotal role of BM vascular dynamics and miR-126 in treatment resistance but also can potentially lead to approaches that enhance the efficacy of FLT3-targeted therapies to eradicate LSCs and improve long-term outcomes for AML.

## Methods

### Agent Based Model of the AML Bone Marrow Microenvironment

The ABM was built on the Hybrid Cellular Automata (HCA) paradigm applied to study evolutionary dynamics and cellular interactions in various biological contexts [16-23]. An HCA framework is defined by discrete agent types updated at each time step based on flowcharts and partial differential equations describing the dynamics of signaling molecules in the microenvironment. The model was implemented using the Hybrid Automata Library (HAL) [24]. The model is defined on a 2D rectangular grid (100 × 100 pixels) representing a section of the AML bone marrow niche, where each pixel represents 10μm^2^ and each timestep represents one 24-hour period. The model has four types of agents, representing blast cells, LSCs, and ECs representing vasculature. The model contains four diffusible molecules, miR-126, TNF-α, a miR-126 inhibitor, miRisten, and an FLT3 TKI, AC220.

### Model Initial Conditions and Agent Rules

When the model is initialized, all cells are placed on the grid randomly, except for the ECs, which are structured based on experimental data. For each timestep, one cell is chosen randomly to perform its actions (Fig. 1). The behavior for each cell is as follows: LSCs undergo a decision process starting with assessing viability. If an LSCs dies, it is removed from the system. If the cell survives and does not divide, it is assumed to re-enter the cell cycle, maintaining its LSC state. However, if an LSC divides, then the LSC either produces another LSC or an AML blast cell dependent upon the current population size of LSCs. If the number of LSCs has dropped below 5% of the domain, which is the initial population size, then an LSC daughter cell is produced. Otherwise, an AML blast daughter cell is produced. This allows the model to capture the behavior in which LSCs can divide symmetrically or asymmetrically [25, 26].

**Figure 1.**
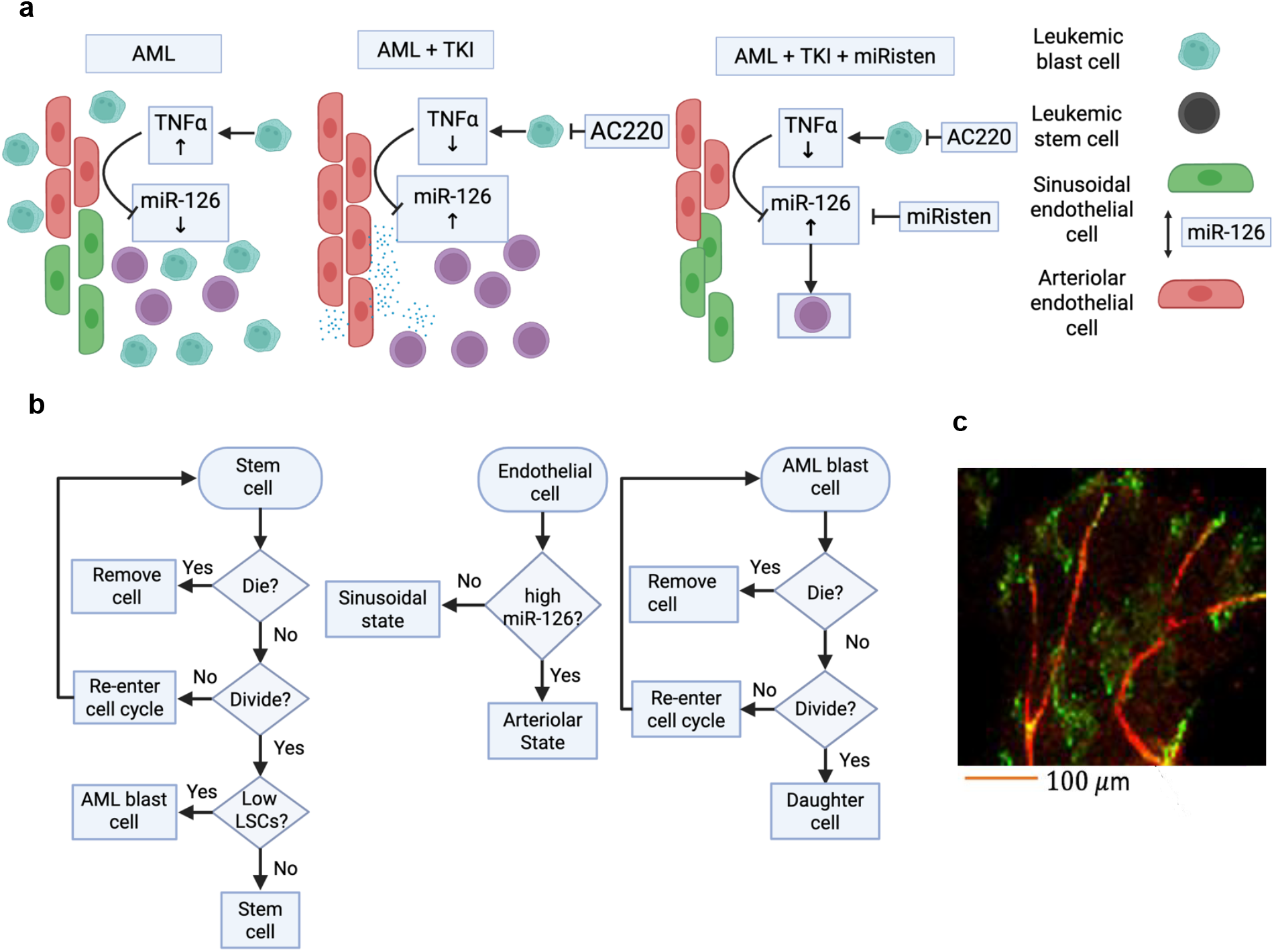
**a** Description of the Janus phenomena. **b** The flowcharts describe the sequence of actions taken by stem cells, blast cells, and endothelial cells. **c** Vascular data from a mouse tibia was used to construct the vascular architectures in the model.

ECs are assumed to transition between two distinct states based on the presence of miR-126. If the local miR-126 concentration is higher than the TNF-α, the EC adopts a miR-126 producing arteriolar state. Conversely, if the miR-126 concentration is lower than the TNF-α concentration, the cell assumes a sinusoidal state, where it does not produce miR-126. This binary state reflects the ECs adaptability and its potential involvement in vascular remodeling and microenvironment regulation. AML blasts follow a similar decision-making pathway. If an AML blast cell dies, it is removed. If it does not divide, it re-enters the cell cycle, remaining as an AML blast. Upon division, blasts cells produce blast cells, contributing to the expansion of the blast population. This behavior highlights the proliferative nature of AML blasts and their role in disease progression.

### ABM Model Parameters

Initial population fractions, growth and death rates for the blast and LSCs were informed from the literature (Supplementary Table 1) [27]. Blast cells can divide once per 24-hour period if there is enough space, producing one blast daughter cell. There is a probability of 0.15 per day that they will die due to naturally occurring apoptosis. This number was empirically derived to ensure the model output (the blast cell count) matches what is observed in the literature. Cell proliferation is implemented as a Bernoulli process, in which each cell has a fixed probability of undergoing division during a given timestep. Specifically, a uniformly distributed random number U(0,1) is drawn each timestep for each cell, and death occurs if this value is less than a specified probability *p*. Here, we assign *p*=0.15, corresponding to a 15% chance of death per timestep. This approach effectively samples from a binomial Bernoulli distribution with success probability *p* (a failure probability of 1-*p*) and allows for probabilistic modeling of asynchronous cell death at the single-cell level. The choice of *p*=0.15 was informed through model calibration, with this number corresponding to an average death rate in the model consistent with observed tumor growth dynamics.

When TNF-α is present locally, blast cells uptake AC220 (a representative TKI) and the probability of death increases proportional to the amount of AC220 [28]. LSCs also can divide once per 24-hour period, with the caveat that they can divide symmetrically or asymmetrically, producing either another LSC or another blast cell. LSCs also have a small probability of natural apoptosis, and AC220 increases the probability of death unless the local concentration of miR-126 is higher than that of AC220, in which case there is no increase in death probability from AC220. The initial population fractions were informed from experimental data, and the model is initialized with 90% of the domain containing blast cells, 5% of the domain consisting of LSCS, and a maximum of 2% of the domain consisting of ECs for any given vascular architecture, informed from experimental data [11]. Treatment induced cytotoxic effects of AC220 on blast cells and LSCs were inferred computationally through comparing model outputs with published data. More specifically, the basal death rates of the blast cells and LSCs is increased based on the concentration of AC220 taken up by each cell. The suppressive effects of miRisten on miR-126 by EC cells was done in a similar manner [11, 27, 28].

### Vascular Architecture and Diffusible Molecule Dynamics

The ABM was initially investigated with a generic vascular structure, and then 16 additional vascular structures were considered, informed by experimental data (Fig. 1c). The vasculature was assumed to be fixed, and there was no modeling of angiogenesis. Both drugs, AC220 and miRisten, diffused through the domain originating from the vasculature, according to a set of partial differential equations (PDEs) (Eq 1). While the vasculature was fixed, the vasculature structure could exist in one of two states, sinusoidal or arteriolar based on the local levels of TNF-α and miR-126, respectively. TNF-α was expressed and diffused from blast cells, while miR-126 diffused from the ECs. Additionally, local levels of miR-126 also impacted the LSC compartment. When the local area around LSC contained an amount of miR-126 higher than that of the base TNF-α concentration, LSCs had decreased sensitivity to AC220, resulting in little to no death of the LSCs due to AC220. For each vascular architecture and treatment condition, 50 simulations each were ran.

TNF-α and miR-126 are produced by the blast cells and ECs respectively at a basal rate of 0.20 μM^2^ per timestep for miR-126 and 0.15 μM^2^ per timestep for TNF-α. These values were inferred computationally based on the molecular weight of the molecules and how quickly they could move through a space the size of the model domain in one 24-hour period. We assume that blasts and ECs produce TNF-α and miR-126 respectively at constant rates, unless miR-126 is inhibited by TNF-α or miRisten. AC220 and miRisten are released from the vasculature at a fixed rate once per day. The concentration of AC220, miRisten, miR-126, and TNF-α are produced, decayed, and passively transported (diffused) through the domain according to the PDE Eq. 1:

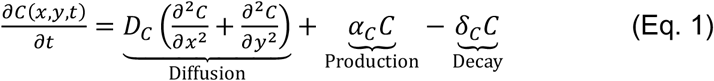

where *C* is the concentration of the diffusible, *D* is the diffusion coefficient, α_*C*_ is the basal production of the molecule, and *δ*_*C*_ is the clearance, or decay, of the molecule. Parameter values were estimated based on the molecular weight of each diffusible as well as approximating the distance the molecule would travel through a space similar in size to the domain of the model in one 24-hour period, the size of the timestep used in the model (Supplementary Table 1). The system of PDEs was solved using the forward time and space-centered space scheme, with periodic boundary conditions in HAL.

## Results

### The ABM Recapitulates the Janus Phenomenon in AML

Once calibrated, the ABM was first tested to determine its ability to recapitulate the Janus phenomena (Fig. 2a-b) within an arbitrarily designed vascular architecture intended to approximate generic blood vessel shapes in a two-dimensional space, with the only constraint that it must fit in the domain of the model. The ABM showed declined of FLT3-ITD blasts once TKI treatment with AC220 began, rapidly followed by reduction of blast-secreted TNF-α and in turn increased EC miR-126 levels. This led to LSCs expansion and eventually to blast re-growth once treatment ended. We observed that the EC populations shifted from a sinusoidal state characterized by miR-126^low^ ECs when TNF-α level was high, to an arteriolar state characterized by miR-126^high^ ECs when TNF-α level dropped due to blast cytoreduction. Thus, the ABM successfully recapitulated the Janus phenomenon, demonstrating that treatment-induced blast cytoreduction resulted in TNF-α decrease and in turn in increased EC miR-126 production and enhanced density of arterioles that supply EC miR-126 to LSCs, which in turn expanded and drove leukemia re-growth (Fig. 2b).

**Figure 2.**
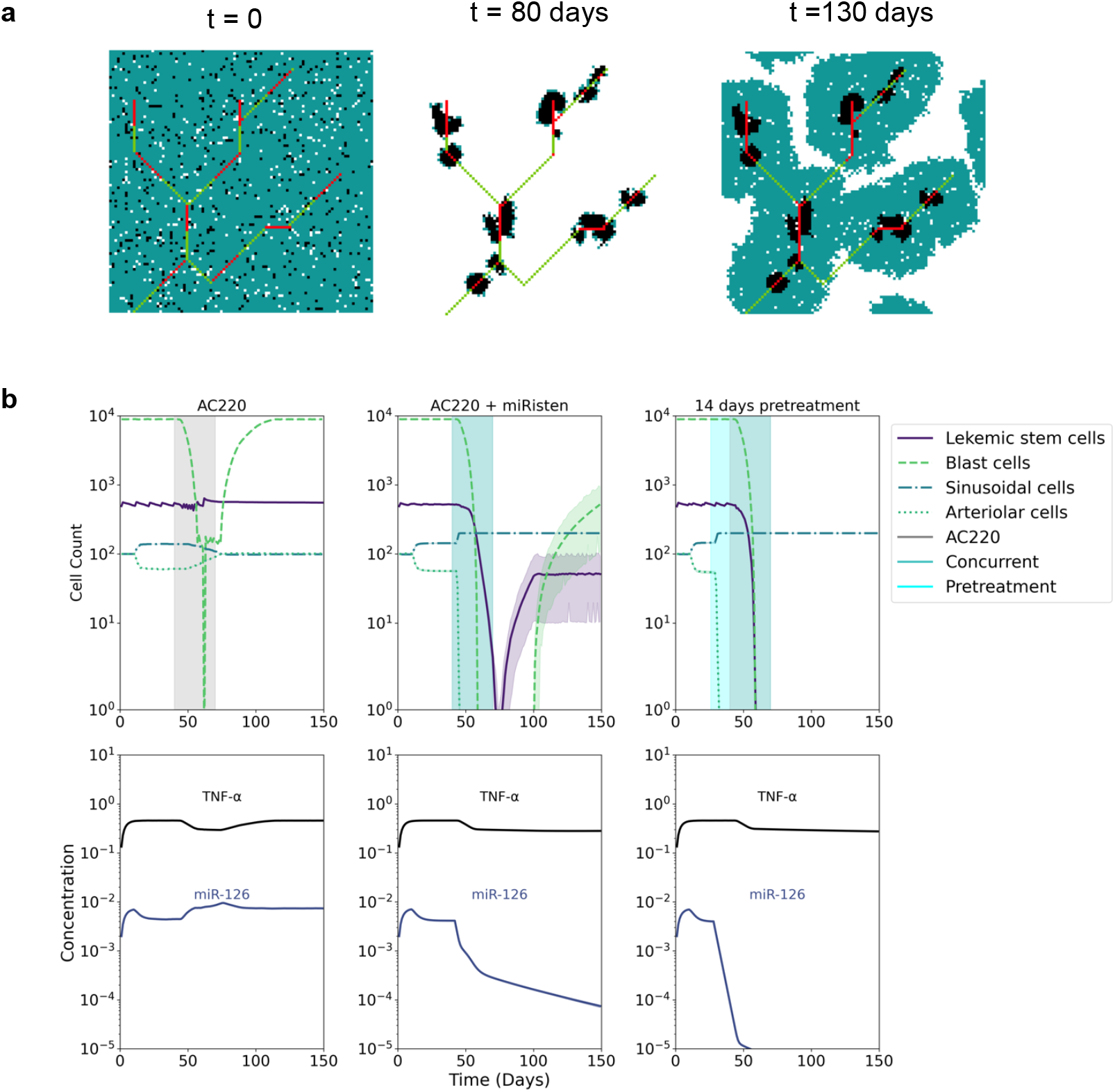
**a** Model visualizations taken from three timepoints. Cell population, TNF-α, and miR-126 results for the base vascular grid for three treatment conditions: **b** AC220 for 30 days, **c** AC220 + miRisten concurrently for 30 days, and **d** 14 days miRisten pretreatment followed by concurrent AC220 and miRsiten for 30 days.

### The Janus Phenomenon can be overcome with different treatment schedules

Next, we analyzed if the Janus phenomenon could be mitigated with administration of miRisten, a novel oligonucleotide that inhibits mature miR-126 [10-12, 14, 15]. When given simultaneously with AC220 (20 mg/kg/day), miRisten (10 mg/kg/day) decreased but did not completely eliminate disease relapse (Fig. 2b). However, the model predicted that administration of miRisten alone for 14 days prior to the start of a treatment with AC2202 alone or combination of AC220/miRisten could prevent disease relapse and progression.

With this data at hand and to determine the optimal duration for miRisten pretreatment, we next tested the ABM with various schedule of miRisten pretreatment given 1 to 14 days before initiation of AC220 alone or AC220/miRisten (Fig. 3b). We observed that miRisten pretreatment, regardless of duration, was effective in reducing relapse and disease progression compared to AC220 or AC220/miRisten given without miR-126 pretreatment. When the total duration of miRisten pretreatment was limited to 30 days, a common duration of a treatment cycle for AML patients [28], a minimum of 2 days of miRisten pretreatment was required to prevent relapse if AC220 was then administered sequentially (Fig. 4b).

**Figure 3.**
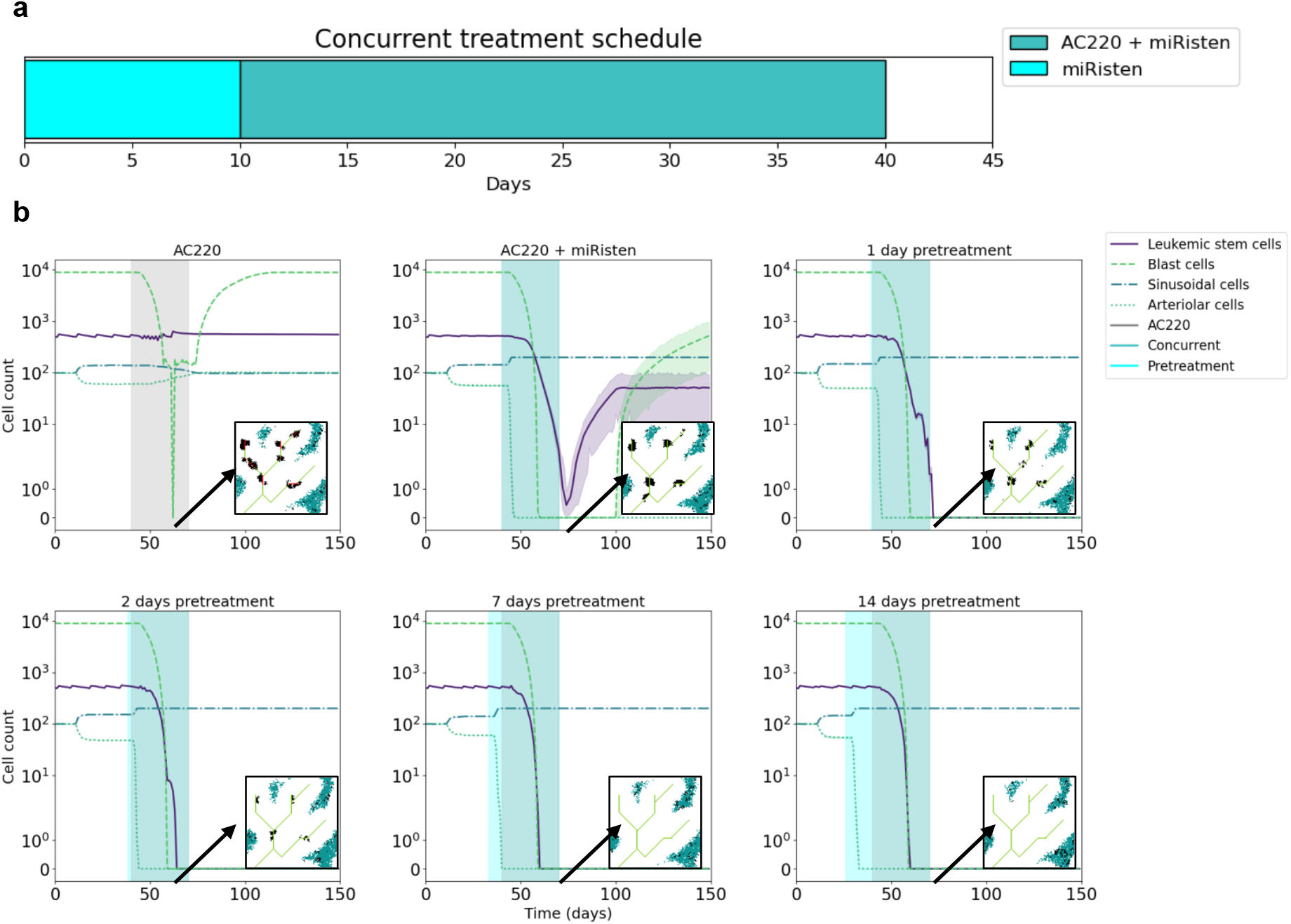
**a** Treatment scheme for pretreatment with miRisten followed by combination AC220 and miRisten. b Results for the base vascular architecture for various combination periods. Inserts are taken at day 55, with AC220 starting at day 40 after the described pretreatment period.

**Figure 4.**
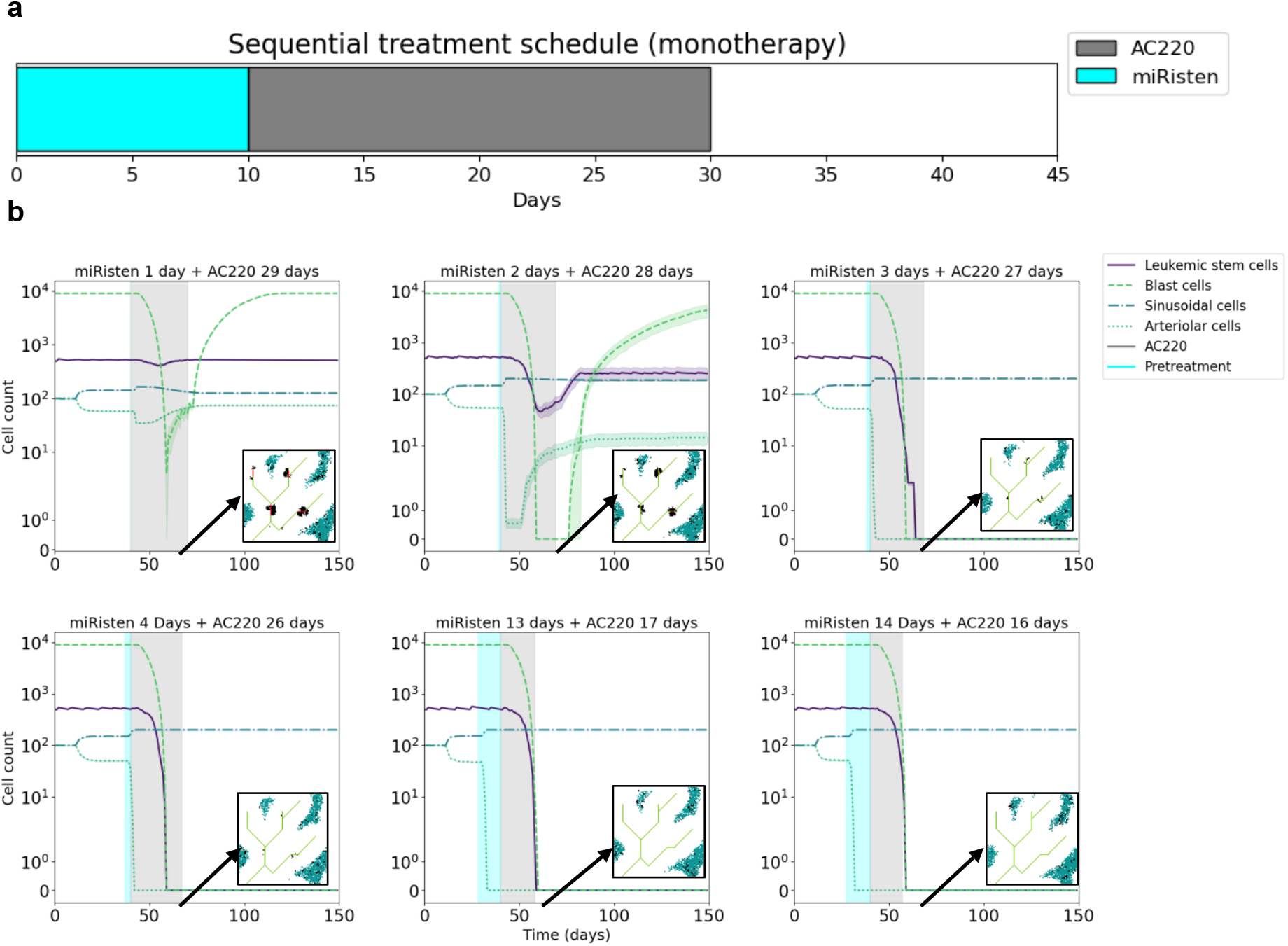
**a** Treatment scheme for pretreatment with miRisten followed by only AC220. b Results for the base vascular architecture for various sequential monotherapy periods. Inserts are taken at day 55, with AC220 starting at day 40 after the described pretreatment period.

### Treatment outcome depends on bone marrow vascular architecture

To connect our ABM model to experimental data, we designed 16 different vascular architecture patterns as informed from the *in vivo* data (Fig. 5a-b). For these 16 different vascular patterns, we tested the activity of pretreatment duration of miRisten followed by a combination of AC220 or miRisten/AC220 (Fig 5c). We found that for most vascular architectures, administering miRisten pretreatment followed by AC220 was more effective than miRisten pretreatment followed by AC220/miRisten, if the ultimate goal was to minimize total drug usage. However, when miRisten dosage was not considered as a limiting factor, the antileukemic activity was similar for miRisten pretreatment followed by AC220 alone or miRisten/AC220. It should also be noted that even though models with individual architectures were cured at different days, there was a single time point at which cure was achieved regardless of the vascular architecture. For the treatment regimen consisting of miRisten pretreatment followed by miRisten/AC220, cure was achieved with 2 days of miRisten pretreatment except for vascular architectures 4 and 8, where cure could be achieved with 1 day of miRisten pretreatment (Fig. 5c). Additionally, we noticed that the different vascular structures primarily fell into one of two groups, either lower ECs and slower response, or a higher number of ECs and a faster response (Fig. 5c). This is likely due to the number of ECs directly correlating to the amount of vasculature within the domain, with a more vascular domain allowing for a greater amount of both miRisten and AC220 to reach the entire domain. For the sequential regimen, when pretreatment was applied followed by only AC220, cure was achieved with as little as 1 day of miRisten pretreatment for all vascular architectures except for 6, which reached cure with 3 days of miRisten pretreatment (Fig. 5c).

**Figure 5.**
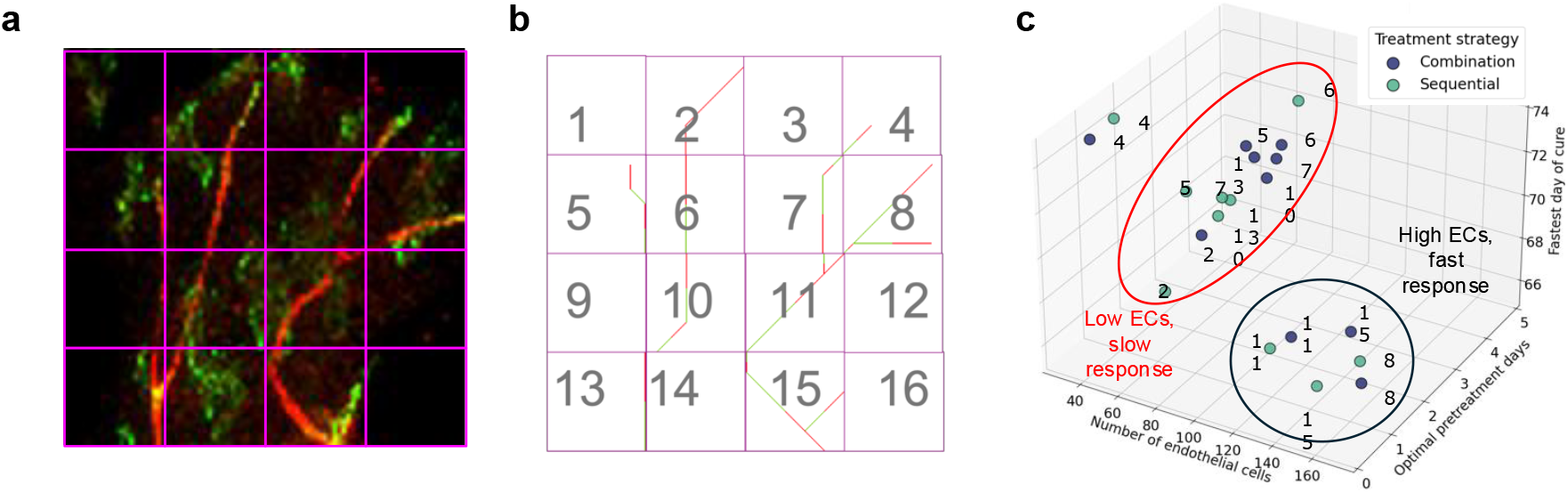
**a** A section of mouse tibia was divided into 16 equal squares and then used to inform the vascular architectures the model. **b** The vascular architecture designs in the model. **c** 3D plot showing the relationship between number of ECs, optimal cure period, and fastest cure period.

### Distance of LSCs from vessels impact on disease response

Noticing that there was a difference in response for some of the vascular architectures to miRisten pretreatment followed by AC220 or miRisten/AC220, we sought to explore if the spatial configuration of the vascular architectures could impact on LSC resistance in the respective BM niches. Thus, we counted the mean number of surviving LSCs based off their distance from the vasculature at the final timestep in the model (Fig. 6a-d, Supplemental Figs. 1-2), where a clone referred to cells derived from a single LSC. All progeny cells were computationally tracked over time, allowing for analysis of spatial dynamics, such as their proximity to vasculature and survival outcomes under different treatment conditions. We observed that for most vascular architectures, pretreatment with miRisten for 1 day followed by 29 days of AC220 significantly reduced the number of LSCs through miR-126 inhibition by miRisten removing the protective vascular niche (Fig. 6a-d, Supplemental figure 1-2). With this data at hand, we then focused on the spatial distribution of surviving clones (Fig. 6a-d, Supplemental Figs. 1-2). We also observed that in vascular configuration #15, all surviving clones were situated relatively close to the vessels (less than 25μm^2^), according to the hypothesis that LSC homeostasis is supported by local vascular miR-126 supply [10, 11]. Of note, for a system configuration with no vasculature, it was impossible to achieve cure as no drug could be delivered.

**Figure 6.**
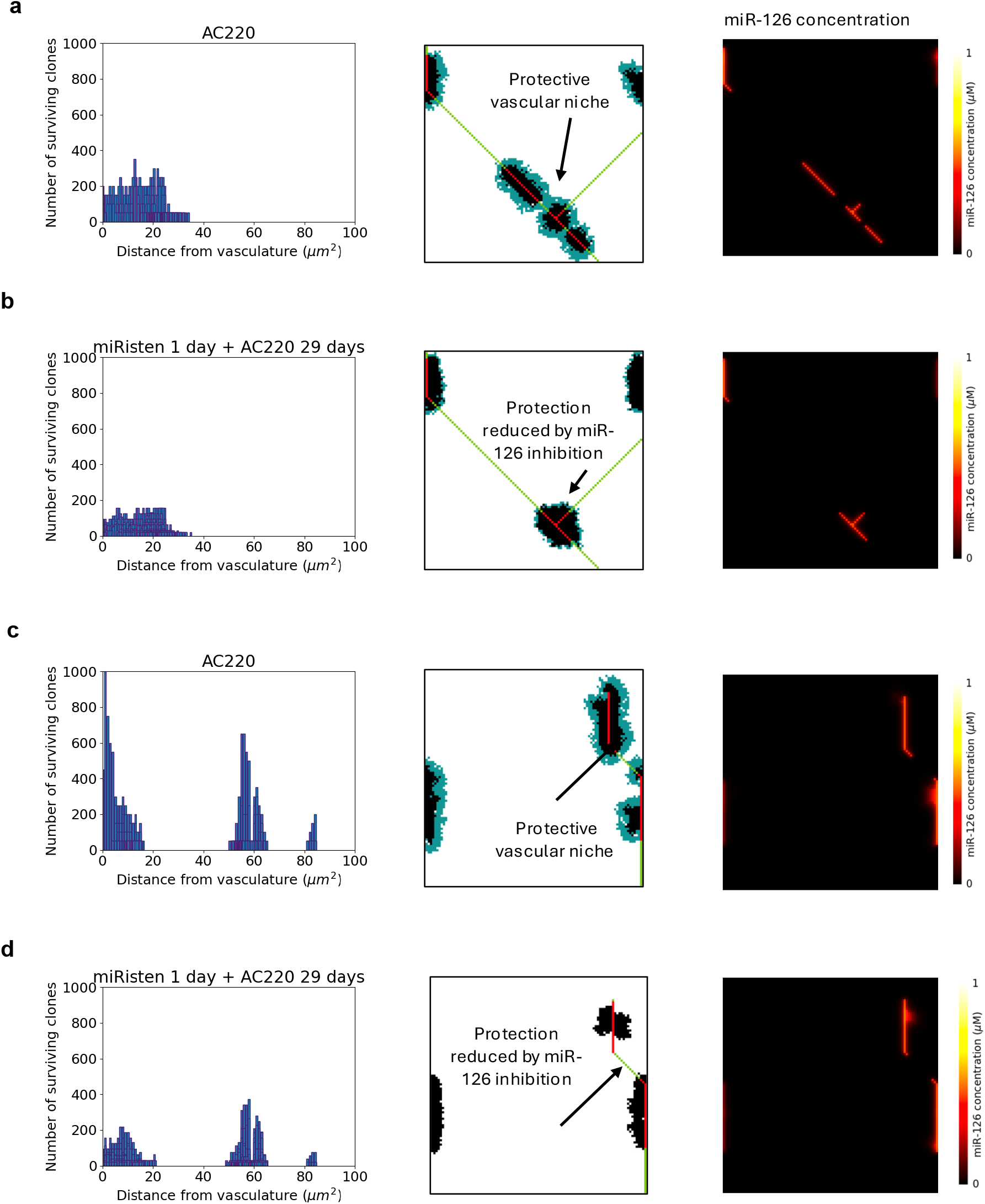
**a** Vascular architecture #15 at timepoint 80, 10 days after treatment AC220 alone. **b** Vascular architecture #15 at timepoint 80, 10 days after 1 day miRisten pretreatment and 29 days of AC220. **c** Vascular architecture #5 at timepoint 80, 10 days after treatment AC220 alone. **d** Vascular architecture #5 at timepoint 80, 10 days after 1 day miRisten pretreatment and 29 days of AC220.

We observed that for all vascular configurations when miRisten pretreatment was followed by AC220, there were less surviving LSCs closer to the vessels than those surviving when no miRisten pretreatment was given and AC220 alone was administered (Fig. 6a-d, Supplemental Fig.1-2a-l). This observation supports the hypothesis that LSC distance from vasculature is a critical determinant of therapy activity. However, we identified a specific treatment timepoint where miRisten pretreatment significantly reduced the Janus phenomenon. This suggests that precise timing of miRisten administration can help mitigate adverse spatial effects on LSC response.

## Discussion

Herein, we demonstrated that an ABM of the BM microenvironment calibrated with experimental data recapitulated a Janus phenomenon in AML [12]. Initial findings revealed that EC miR-126 production increased upon administration of cytotoxic therapy due to depletion of TNF-α producing myeloblasts. Decrease in TNF-α resulted in a surge in EC miR-126 production, increased arteriolar vascularization and enhanced EC miR-126 supply to LSCs, which in turn retracts in quiescence and become invulnerable to treatment, posing the basis for future disease relapse. Preventing EC miR-126 production with miR-126 inhibition with miRisten mitigated the paradoxical remodeling of the BM vasculatures that during cytotoxic treatment provides LSC protection that we called Janus phenomenon. Pretreatment with miRisten followed by treatment with AC220 or miRisten/AC220 captured optimal dose schedules that prevented disease relapse while minimizing the amount of drugs given. This model conceptually validates our previous experimental findings and underscores the usefulness of ABMs in understanding how to mitigate optimize drug schedule and delivery in AML [10-12].

Analysis of 16 different vascular patterns informed by experimental data revealed that pretreatment with miRisten followed by either concurrent miRisten/AC220 or AC220 alone, prevented relapse under specific conditions, although not equivalent. Spatial analysis of LSC survival revealed the role of vascular structures in protecting LSCs from the cytotoxic effects of therapy (Fig. 6a-d). While there is no exact minimal distance from the vasculature that shelters the LSCs across all vascular architectures in the model, we did notice that the closer to the vessel the LSCs were located, the more the protective effect of EC miR-126.

These insights emphasize the spatial dependency of therapeutic efficacy, where proximity to vasculature significantly affects the potential success of anticancer therapy in eradicating AML. Thus, our study provides a computational framework for understanding how vascular architectures and treatment timing synergistically impact outcomes that matches the experimental observation and allows for exploration of other therapeutic strategies that target the complex interplay between BM microenvironment and LSCs. Of note, while it is impossible to know the precise vascular architectures in patients, herein we show that there can exist a minimum number of days for miRisten pretreatment that could mitigate the Janus phenomena for most vascular architectures in the BM microenvironment. This could pave the way for future advances in AML personalized treatment, where geometrical and structural features of the BM niche could determine the type and schedule of treatment. In fact, beyond AML and TKIs, our findings may have broader therapeutic implications for other hematologic malignancies that exhibit stem cell persistence and microenvironment-mediated resistance, as well as other treatment modalities such as BCL-2 inhibitors [29] or immune-based interventions where microenvironment-driven resistance may play a significant role. Similar modeling approaches could also be extended to solid tumors where the vascular niche may play a similar protective role in cancer stem cells’ survival.

A key strength of this study is the integration of computational modeling with experimental vascular architectures, allowing for hypothesis-driven exploration of combination treatment strategies *in silico*. The ABM enables spatially explicit simulations that capture heterogeneity in therapeutic response due to vascular proximity, a factor difficult to assess in traditional preclinical models. This methodology could be extended to study other microenvironmental interactions, such as stromal-mediated drug resistance or immune evasion mechanisms in cancer. However, like all computational models, our approach has limitations. The ABM is constrained by predefined assumptions regarding cell behavior, vascular dynamics, and the production, decay, and diffusion of molecules in the tissue. Additionally, while our vascular structures were informed by experimental data, they are two-dimensional simplifications of complex and heterogeneous three-dimensional tissues. Future refinements could incorporate patient-derived imaging data to personalize model predictions and improve clinical translatability.

Nevertheless, our findings highlight the importance of vascular architecture in shaping AML treatment responses and propose a strategy for mitigating the Janus phenomenon through miRisten pretreatment. By integrating computational modeling with experimental data, this study provides a foundation for optimizing treatment timing and microenvironment-targeted therapies. Further experimental and clinical validation will be essential to translate these insights into patient care, potentially improving outcomes for AML patients.

