## Supplemental Information for "Overcoming treatment resistance mediated by the bone marrow vascular niche in acute myeloid leukemia"

### Supporting Information Text

**Methods:** Random seeds were used for reproducibility so that saving large datasets were not required to make results reproducible. Code can be found here: https://github.com/mfroid/JanusABM

Supplemental 1

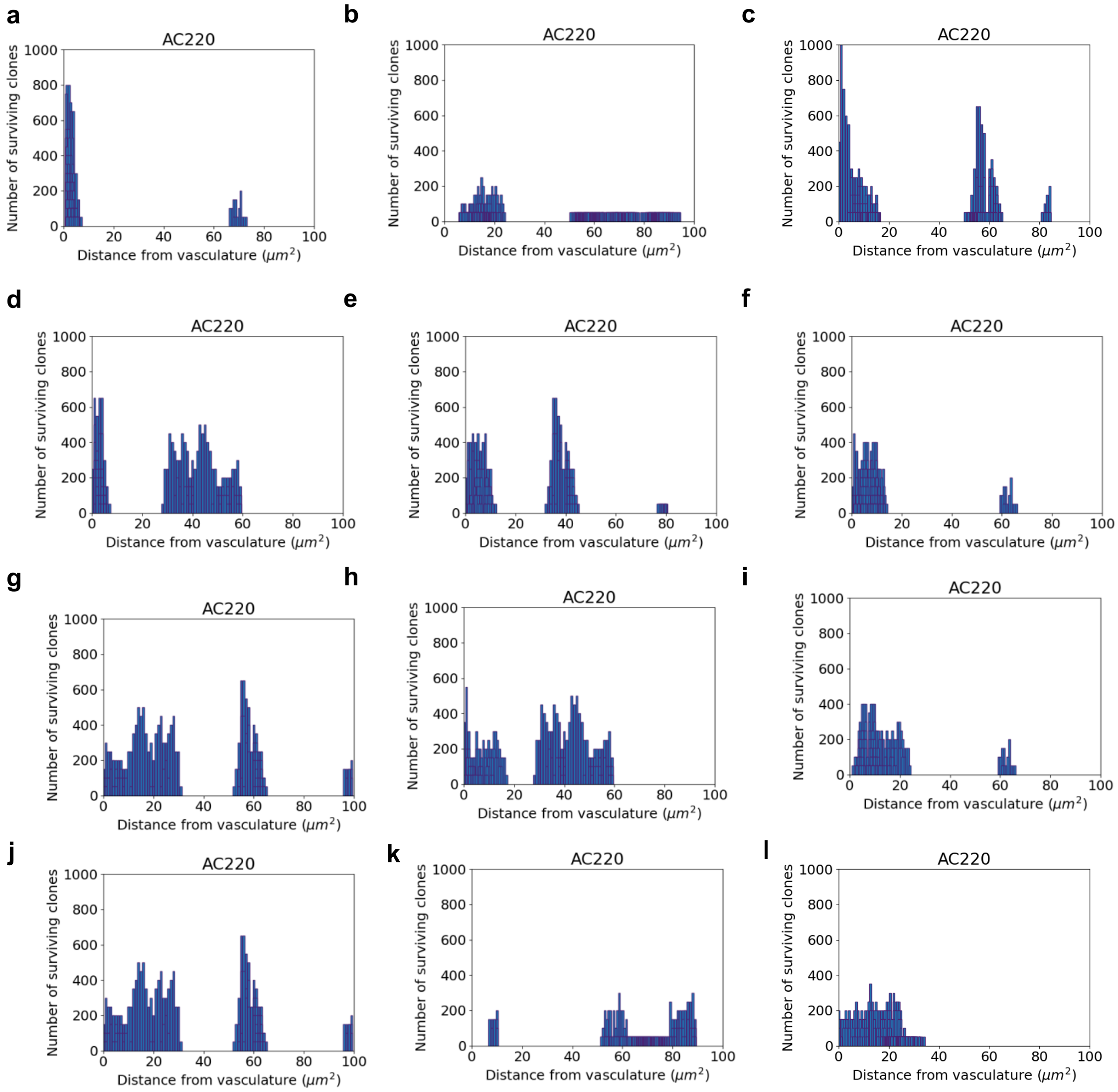


**Fig. S1. a-l** Surviving clones for all vascular architectures under AC220 only.

Supplemental 2

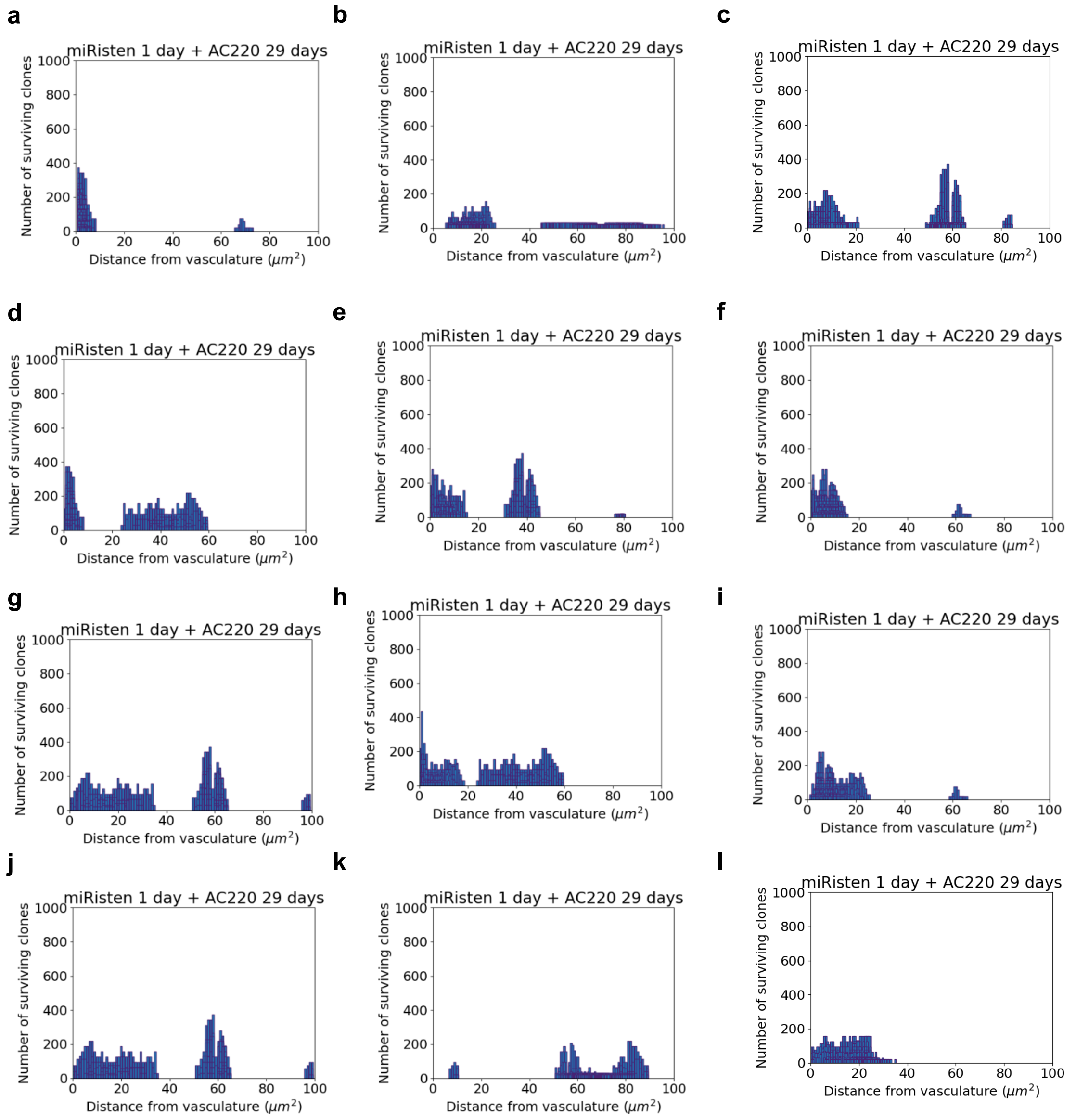


**Fig. S2. a-l** Surviving clones for all vascular architectures after 1 day miRisten pretreatment followed by AC220 only.

| Parameter | Value | Source |
|---|---|---|
| Blast cell growth rate | $0.70$ day$^{-1}$ | 1 |
| Stem cell growth rate | $0.50$ day$^{-1}$ | 1 |
| Blast cell death rate | $0.15$ day$^{-1}$ | 1 |
| Stem cell death rate | $0.015$ day$^{-1}$ | 1 |
| miR-126 diffusion rate | $0.2$ $\mu$m$^2$/day | Inferred based on molecular weight to allow for degradation within the model timestep that allowed model outputs to match empirical observation (1, 2). |
| miR-126 decay rate | $0.01$ day$^{-1}$ | Inferred to allow for degradation within the model timestep that allowed model outputs to match published empirical data (1, 2). |
| TNF$\alpha$ diffusion rate | $0.1$ $\mu$m$^2$/day | Inferred considering TNF$\alpha$'s molecular weight and expected range of paracrine signaling in tissue and so that degradation within the model timestep allowed model outputs to match published empirical data (1, 2). |
| TNF$\alpha$ decay rate | $0.01$ day$^{-1}$ | Inferred based on molecular weight to allow for degradation within the model timestep that allowed model outputs to match published empirical data (1, 2). |
| AC220 diffusion rate | $2.5$ $\mu$m$^2$/day | Inferred based on the molecular size of the drug and the need for plausible spread across the domain in each 24-hour timestep. |
| AC220 decay rate | $0.01$ day$^{-1}$ | Inferred based on molecular weight to allow for degradation within the model timestep that allowed model outputs to match published empirical data (1, 2). |
| miRisten diffusion rate | $0.133$ $\mu$m$^2$/day | Inferred based on the molecular size of the drug and the need for plausible spread across the domain in each 24-hour timestep (1, 2). |
| miRisten decay rate | $0.01$ day$^{-1}$ | Inferred based on molecular weight to allow for degradation within the model timestep that allowed model outputs to match published empirical data (1, 2). |

1

**Table S1.** Table showing model parameters.
